# Nipah virus matrix protein utilizes cortical actin to stabilize the virus assembly sites and promote budding

**DOI:** 10.1101/2025.01.27.635114

**Authors:** Jingjing Wang, Vicky Kliemke, Jinxin Liu, Giuliana Leonarda Matta, Qian Wang, Yuhang Luo, Mengyu Zhang, GuanQun Liu, Qian Liu

## Abstract

Several families of enveloped viruses assemble and bud from the host cell plasma membranes (PM), including paramyxoviruses. Nipah virus (NiV) is a deadly zoonotic paramyxovirus causing yearly outbreaks in Southeast Asia with >75% mortality. NiV encodes matrix proteins (M) that drive assembly and budding. NiV-M forms dimers and interacts with membrane lipids in the host cell’s PM for budding. Using single-molecule localization microscopy and single-particle tracking, we show that the host F-actin maintains the nanoscale organization of NiV assembly sites at the PM. This F-actin-dependent integrity of NiV assembly sites is observed at the membrane retention stage after the assembly process is complete, rather than during the recruitment of NiV-M molecules to these sites. NiV-M interacts with actin via its C-terminal domain. We also show that the actin-branching factor, Arp2/3 complex, promotes virus-like-particle production. Meanwhile, inhibiting Arp2/3 disfavors the PM retention of NiV assembly sites and NiV-M-mediated generation of membrane protrusions, but does not affect the assembly rate. This suggests that Arp2/3-nucleated actin polymerization and branching are critical for maintaining NiV-M assembly sites at the PM, allowing it to facilitate membrane protrusion generation for virus budding. Our findings support the following model: NiV-M interacts with F-actin through its C-terminus to remain at the host PM after assembly completion. This F-actin-dependent retention is promoted by the Arp2/3-driven actin branching and polymerization, which also drives the formation of NiV-M-induced membrane protrusions necessary for virus budding.

**Significance:** Nipah virus (NiV) is a deadly paramyxovirus capable of animal-animal and animal-human transmissions. To produce a NiV particle, the NiV matrix protein (M) must create an assembly site by binding to the surface of an infected cell and pushing the cell membrane outward for virus budding. To do so, the NiV matrix protein must co-opt or overcome an actin network underneath the cell membrane. We provide single-molecule evidence that M interacts with the actin cytoskeleton for membrane retention but not recruitment of M to existing assembly sites. This process is promoted by the Arp2/3-driven actin branching and polymerization. Our findings suggest the role of actin remodeling in NiV budding and identify a druggable site at the C-terminus of NiV-M.

## Introduction

Paramyxoviruses are a group of clinically important viruses which are mainly transmitted via respiratory droplets and direct contact (1, 2). Measles (MeV) and human parainfluenza viruses (PIV) are established human pathogens and cause infections in all age groups and hospitalization every year (3, 4). Nipah (NiV) and Hendra (HeV) viruses in the *henipavirus* genus are known zoonotic viruses that cause yearly outbreaks in southeast Asia and Australia with over 70% mortality rate in humans (5, 6). They are capable of animal-animal, animal-human, and human-human transmissions (5, 6). Despite these threats, no vaccines or therapeutics are approved for humans against these emerging paramyxoviruses (6).

Paramyxoviruses are enveloped, negative-strand, non-segmented RNA viruses (1). The RNA genome codes for six proteins, including two membrane glycoproteins for virus entry—the fusion protein (F) and the attachment protein (H/HN/G), the matrix protein (M), and three proteins for ribonucleoprotein (RNP) complex formation and RNA genome replication—the nucleoprotein (N), the large polymerase protein (L), and the phosphoprotein (P) (1). Among them, M orchestrates the assembly and budding of most paramyxoviruses and thus plays a central role in virus transmission among host cells (1). A cryoelectron tomography (cryo-ET) study reveals that MeV-M protein forms two-dimensional (2D) arrays on the inner leaflet of the plasma membranes (PM) at the MeV assembly and budding sites in infected cells and cell-free virus particles, similar to that of Newcastle Disease viruses (NDV) (7, 8). MeV-M proteins have also been observed to coat RNPs but dissociate with the viral membranes (9). X-ray crystallography has shown that the paramyxovirus M forms dimers and associates with PM by interacting with membrane lipids, phosphatidylserine (PS), and phosphatidylinositol 4,5-biphosphate [PI(4,5)P_2_], in the inner leaflet (7, 10). Binding to PS and PI(4,5)P_2_ triggers conformational and electrostatic changes in M for membrane curvature generation (10), although the involvement of host factors in this process remains unclear.

The actin cytoskeleton consists of actin filaments (F-actin) organized into higher-order arrays capable of dynamic remodeling (11). Among them, the actin cortex is defined as an actin network underneath the PM (11). Many intracellular pathogens co-opt host cortical actin network for fulfilling the life cycles (12–15). In paramyxoviruses, MeV production is partially inhibited by actin polymerization inhibitors or stabilizers as they affect the intracellular trafficking of RNPs to the PM and virion maturation (16). MeV was observed to bud from microvillus-like structures where the negative-directed actin filaments are tightly associated with the budding virions (17). The M proteins of Sendai virus (SeV) and NDV have been shown to interact with actin filaments directly (18). Further, human immunodeficiency virus-1 (HIV-1) manipulates the actin cytoskeleton for virus production by directly interacting with the F-actin via Gag, and the HIV-1 Gag particle release from CD4 T cells depends on the Rac1–IRSp53–Wave2–Arp2/3 signaling pathway (19). The Arp2/3 complex nucleates the formation of branched actin essential for actin cortex and most filopodia (20, 21). Respiratory syncytial virus (RSV) requires Arp2 for the production of infectious progeny virions, and the Arp2-dependent filopodia formation and cell motility facilitate cell-to-cell virus spread (22). The Arp2/3-mediated actin branching prevents HIV-1 viral particle production in infected T cells, partly correlated with a decreased number of HIV-1 Gag clusters on the T cell membrane (23). Nonetheless, Arp2/3 and cofilin were identified as cellular binding partners of NiV-M in a proteomics-based interaction study (24). We aimed to investigate whether and how NiV-M hijacks cortical actin remodeling for assembly and budding.

Here, we show that NiV-M directly interacts with actin, which is key for maintaining the nanoscale NiV assembly sites at the PM and virus-like-particle (VLP) production. Analysis of assembly kinetics reveals that NiV assembly sites transit through three stages at the PM: assembly, membrane retention, and release. The duration of membrane retention is sensitive to the local F-actin density and NiV-M-actin interaction. Interestingly, the Arp2/3 complex-nucleated actin polymerization and branching facilitate membrane retention of NiV assembly sites and the generation of NiV-M-mediated membrane protrusion. Our data show that NiV-M exploits Arp2/3-mediated actin remodeling to facilitate membrane protrusion generation and viral budding, without affecting virion assembly.

## Results

### NiV-M interacts with actin in the host cell for VLP production

Previous studies show that SeV infection induces actin remodeling to promote efficient virion production by an actin-binding domain at the C-terminal domain of the M protein (18, 25). We aligned the protein sequences of NiV-M, hPIV-1-M, SeV-M, and three actin-binding proteins— WASP, WASF2, and WIPF1. We found homologous residues isoleucine 349 (I349) among all protein sequences and lysine 351 (K351) among all viral protein sequences (Fig. 1A). These two residues are located in the C-terminal domain (CTD; orange) of NiV-M, protrude outward of the NiV-M dimer, and are spatially away from the key motifs for membrane association (cyan) and dimerization (yellow) (Fig. 1B). We mutated I349 and K351 to alanines (A) and fused them to an N-terminal 3xFLAG tag. When expressed in 293T cells, both I349A and K351A pulled down less β-actin than the wild-type (wt) protein, with I349A even less than K351A (Fig. 1C). This suggests that both residues are essential for maintaining the interaction between NiV-M and β-actin. To analyze the role of NiV-M-actin interaction in VLP production, wt and mutant M were ectopically expressed in 293T cells, and the cell lysates and VLPs in the supernatant were analyzed by western blot analysis (Fig. 1D). Normalized budding index shows that I349A and K351A are less efficient in VLP production compared to the wt (Fig. 1D). Next, COS-7 cells were used to examine membrane protrusions induced by NiV-M constructs because these cells have simple membrane structures that facilitate microscopic observations (Fig. 1E) (10). COS-7 cells stably expressing I349A exhibited fewer membrane protrusions compared to cells expressing the wt protein, while no significant difference was observed between K351A- and wt-expressing cells (Fig. 1E). These data indicate that the interaction between NiV-M and F-actin is required for generating membrane protrusions during VLP budding. Additionally, we analyzed the incorporation of β-actin into individual VLPs using confocal microscopy. NiV-VLPs were visualized through GFP fluorescence from GFP-tagged NiV-M constructs (green), while β-actin was labeled using an anti-β-actin antibody (magenta) (Fig. 1F). We confirmed that the GFP-tagged and FLAG-tagged NiV-M constructs showed a similar trend in VLP production (Fig. S1). We detected a higher percentage of β-actin-positive VLPs (GFP+/β-actin+) in the supernatant collected from 293T cells expressing NiV-M-wt compared to those expressing M-I349A and M-K351A, indicating that actin incorporation into VLPs is impaired by these mutations (Fig. 1F). Collectively, these data suggest that the C-terminal actin-binding domain of NiV-M is crucial for actin interaction, VLP production, and actin incorporation into VLPs.

**Fig. 1.**
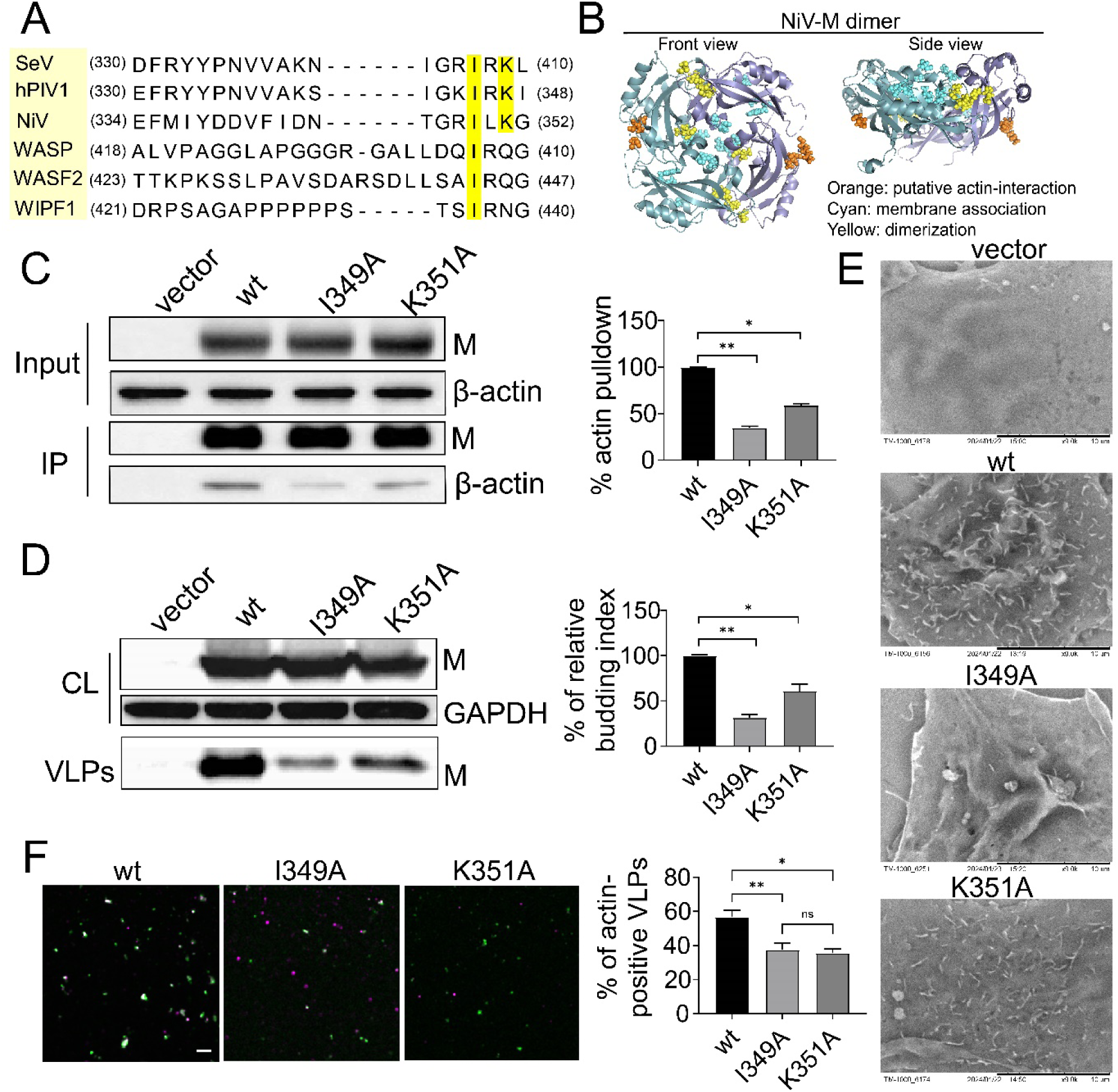
An actin-binding domain in NiV-M is key for NiV-M-actin interaction and NiV VLP production. (**A**) The sequences of M proteins of three paramyxoviruses and three actin-binding proteins are aligned: Sendai virus (SeV), human parainfluenza virus 1 (hPIV1), Nipah virus (NiV), Wiskott-Alderich syndrome protein (WASP), WASP family member 2 (WASF2), and WASP interacting protein 1 (WIPF1). Conserved amino acids are highlighted. (**B**) Key residues for membrane association (cyan) and dimerization (yellow) are away from the actin-binding motif (orange) at the C-terminus of NiV-M. Left, NiV-M dimer crystal structure front view; right, side view. NiV-M structure was modelled using the PyMOL software (https://pymol.org/2/<u>)17</u>. (**C**) Co-immunoprecipitation assay of 3xFLAG-tagged NiV-M constructs and β-actin. 293T cells were transfected with 3×FLAG-tagged NiV-M constructs (wt, I349A, and K351A) along with an empty pcDNA3 vector. At 48 hrs post-transfection, cells were lysed, and β-actin was immunoprecipitated by anti-FLAG magnetic beads. Total cell lysate (Input) and immunoprecipitated proteins (IP) proteins were separated by 10% SDS-PAGE and immunoblotted with mouse α-FLAG (NiV-M detection) or mouse anti-β-actin. Proteins were detected using HRP-conjugated secondary antibodies. Relative actin pulldown is determined based on integrated immunoblot density. Results from at least three independent experiments are shown. (**D**) Western blot of cell lysates and VLPs collected from 293T cells transfected by empty pcDNA3 vector (vector) and plasmids coding for 3×FLAG-tagged NiV-M-wt, I349A, and K351A. GAPDH was used as a loading control. NiV-M was detected using mouse anti-FLAG antibody and GAPDH mouse anti-GAPDH antibody. Proteins were detected by HRP-conjugated secondary antibodies. Relative budding index is determined based on integrated immunoblot density. Results from at least three independent experiments are shown. (**E**) Scanning electron microscopy was used to observe the membrane protrusions of COS-7 cells stably expressing 3×FLAG-tagged NiV-M-wt, I349A, and K351A constructs along with an empty pQCXIP vector. Scale bar:10 μm. Representative images from at least three independent experiments are presented. (**F**) VLPs produced by 293T cells transfected by GFP-tagged NiV-M wt, I349A, and K351A were imaged using laser scanning confocal microscopy. VLPs were purified using ultracentrifugation on a 20% sucrose cushion. Purified VLPs were immobilized on fibronectin-coated cover glass and subjected to imaging. VLPs (green) were localized using GFP and β-actin (magenta) was detected using a mouse anti-β-actin antibody and a donkey anti-mouse antibody conjugated to AlexaFluor® 647. Scale bar: 5 μm. The percent of actin-positive VLPs was determined by the number of VLPs positive for both GFP (M detection) and AlexaFluor® 647 (β-actin detection) over the number of VLPs positive only for GFP (n > 200 for each type of VLPs). Results from at least three independent experiments are shown. Bars represent mean ± SEM (C and D) mean ± SD (F). *p* value was obtained using Welch’s *t*-test. ns: *p* > 0.05, *: *p* ≤ 0.05, **: *p* ≤ 0.01.

### The nano-organization of NiV-M assembly sites is disrupted by F-actin depolymerization

We noticed that NiV-M co-partitioned with vimentin to the cytoskeleton portion of the cell lysate, suggesting that NiV-M is mainly associated with F-actin (Fig. S2A). PK13 cells treated with an F-actin depolymerization drug, Latrunculin A (LatA), were smaller in cell size and showed fewer fine actin structures compared to the vehicle (DMSO) control (Fig. 2A). We then examined the role of F-actin in NiV-M assembly by analyzing the nano organization of NiV-M assembly sites using single-molecule localization microscopy (SMLM). The localization precision of our custom-built SMLM system is ∼10 nm in the lateral direction and ∼25 nm in the axial direction (26, 27). GFP-tagged NiV-M constructs were used to identify NiV-M puncta under widefield microscopy. SMLM images were acquired using Cy3B or AlexaFluor 647 attached to the GFP tag. We verified that the labeling tags at the N-terminus of NiV-M minimally affected NiV-M clustering (Fig. S2B-G). Images show that NiV-M form nanoclusters at the ventral membrane of PK13 cells, consistent with those observed at the dorsal membrane of these cells (Fig. 2B and C) (27). By visual inspection, NiV-M localizations formed distinctive clusters in DMSO-treated PK13 cells (control) and less defined clusters in LatA-treated cells (Fig. 2C and D). To analyze the partition of NiV-M localizations in clusters, NiV-M clusters were identified using a DBSCAN (Density-Based Spatial Clustering of Applications with Noise) algorithm that links closely situated localizations in a propagative manner (26, 28), and the clusters were masked in red in cluster maps (Fig. 2C and D). Smaller clusters and more unclustered localizations (gray) were observed for NiV-M localizations in LatA-treated cells compared to the control (Fig. 2C and D, cluster map). We performed Hopkins’ analysis to evaluate the clustering tendency of NiV-M localizations. The Hopkins’ index indicates the extent of clustering, ranging from 0 (random distribution) to 1 (extreme clustering). The result shows that NiV-M localizations in LatA-treated cells are less likely to cluster compared to the control, demonstrated by a significantly lower Hopkin’s index (Fig. 2E). Consistently, we noticed that a lower percentage of NiV-M localizations segregated into clusters in LatA-treated cells (Fig. 2F). Additionally, NiV-M clusters were smaller upon LatA treatment (Fig. 2G), although they are similarly packed (Fig. 2H). We noticed that the NiV-M clusters in LatA-treated cells less circular (Fig. 2I), consistent with the idea that NiV particles exhibit irregular shapes, as these NiV-M clusters likely serve as sites for virus assembly and budding (27, 29). The total density of the selected regions was similar in both DMSO- and LatA-treated cells, indicating that these differences are not due to varying amounts of NiV-M localizations (Fig. 2J). Taken together, these results suggest that F-actin in host cells promotes NiV-M clustering at the PM, thereby enhancing NiV VLP production.

**Fig. 2.**
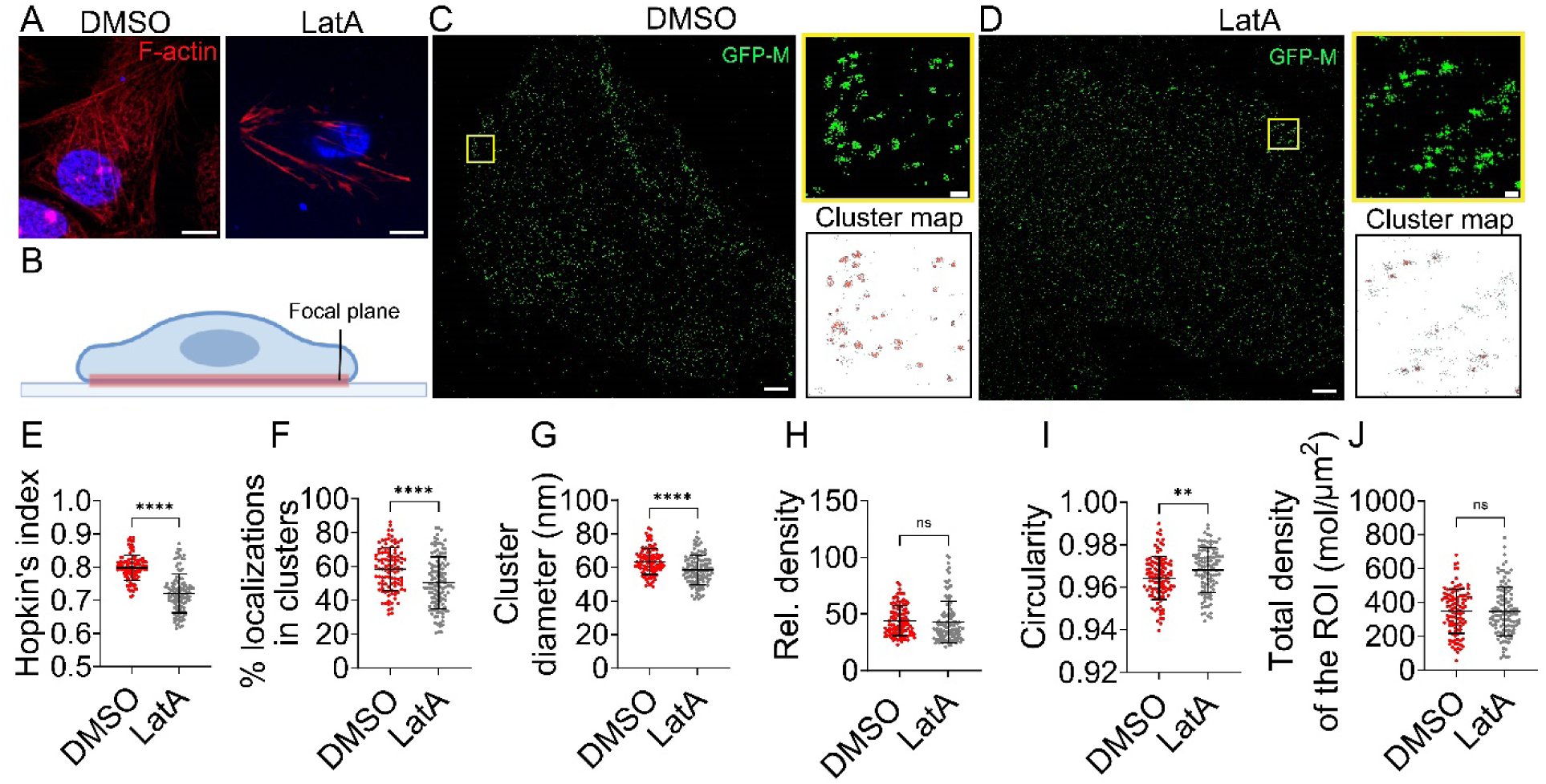
F-actin depolymerization alters the nano-organization of NiV-M. (**A**) PK13 cells were treated with 1 µM DMSO or 1 µM LatA for 5 minutes and stained with phalloidin AlexaFluor^®^ 647. Images were taken using laser scanning confocal microscopy. Representative images of 10-20 cells collected from three independent experiments are shown. Scale bar: 1 μm. (**B**) The Imaging plane is at the ventral membrane of PK13 on the cell-cover glass interface. (**C** and **D**) x-y cross-section (600 nm thick in z) of the SMLM images of GFP-NiV-M at the ventral membrane of PK13 cells treated by DMSO (**C**) or LatA (**D**). The boxed regions are enlarged, and the cluster maps of the NiV-M localizations are shown. NiV-M clusters are masked in red while the unclustered NiV-M localizations are gray. Scale bar: 1 μm and 200 nm. (**E**) The Hopkin’s index of NiV-M localizations in DMSO and LatA-treated cells. The percentage of localizations in clusters (**F**), cluster diameter (**G**), relative density (**H**), circularity (**I**), and total density of the ROI (**J**) are shown in dot plots. Sample size n = 106 (DMSO) and 115 (LatA) from 9-15 cells per group. Bars represent mean ± SD. *p* value was obtained using Welch’s *t*-test. ns: *p* > 0.05, **: *p* ≤ 0.01, ****: *p* ≤ 0.0001.

### The nano-organization of NiV-M assembly sites depends on its proximity to F-actin

To precisely analyze the role of F-actin in NiV-M assembly, we performed dual-color SMLM imaging of NiV-M and actin cytoskeleton on the ventral membrane of PK13 cells. The SMLM images show that NiV-M clusters are distributed around F-actin, with some clusters located close to F-actin and others far away (Fig. 3A-C). To quantitatively compare NiV-M clusters with varying distributions relative to F-actin, we categorized them into two groups based on their association with F-actin: NiV-M clusters that colocalized with F-actin (on) and those that did not (off). First, NiV-M clusters were identified using a DBSCAN algorithm (Fig. 3B-C, cluster map). Next, the clusters were grouped based on their co-clustering with F-actin using a coordinate-based correlation assay for SMLM data (Fig. 3D) (27, 30). The co-clustering between NiV-M and F-actin was mainly observed at the border of the NiV-M clusters, marked by orange-red dots in the correlation map (Fig. 3B and C). Our analysis shows that NiV-M clusters on F-actin are larger (Fig. 3E) and denser (Fig. 3F) than those off F-actin. Interestingly, NiV-M clusters associated with F-actin often displayed irregular shapes, whereas those distant from F-actin were predominantly round (Fig. 3G). These results show that the NiV-M nanoclustering is correlated with its proximity to F-actin.

**Fig. 3.**
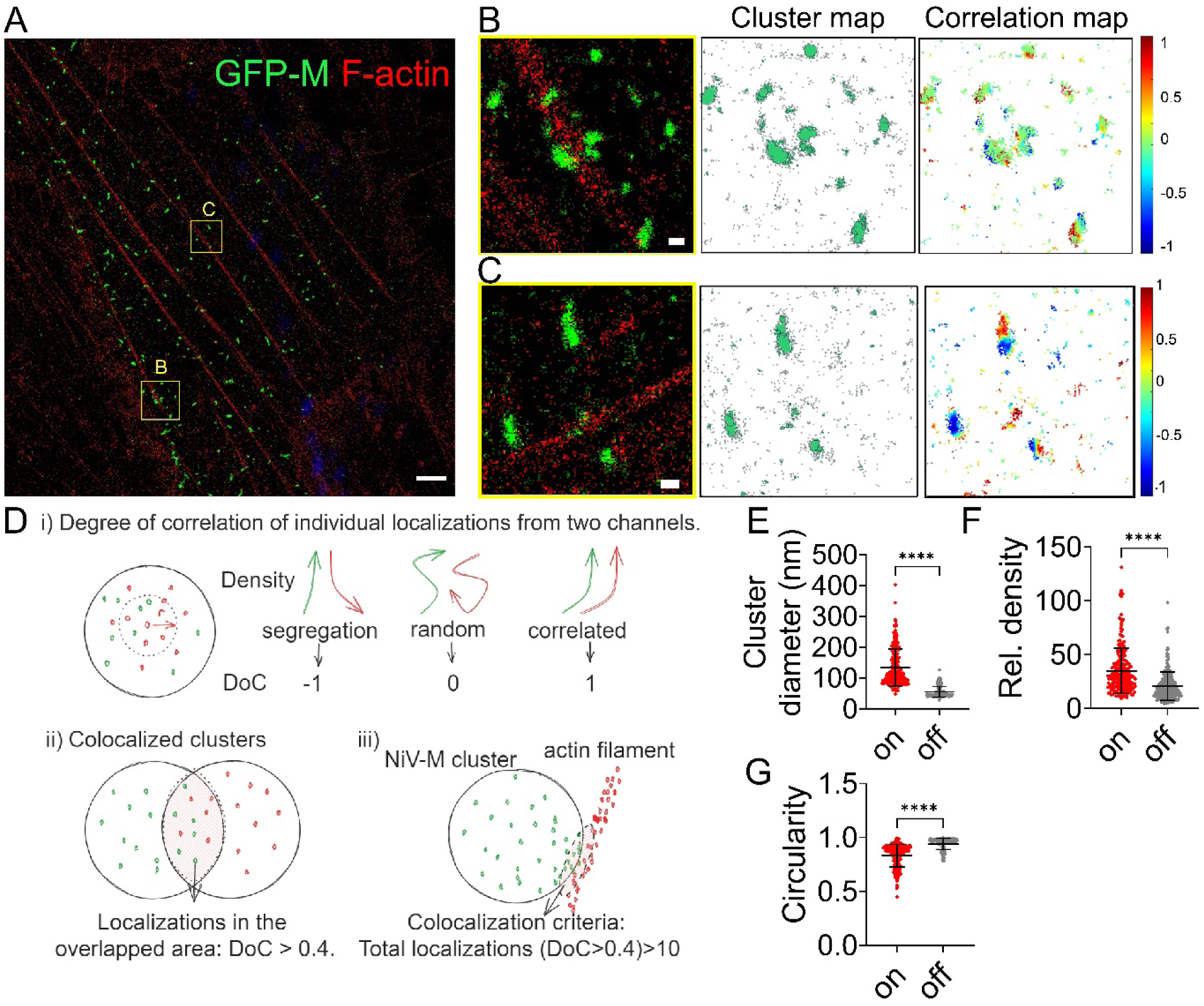
The nano-organization of NiV-M depends on its proximity to F-actin. PK13 cells were transfected with GFP-NiV-M and fixed at 24 hrs post-transfection. PK13 cells were permeabilized and stained with goat anti-GFP antibody and donkey anti-goat antibody conjugated to cy3B (detect M), and phalloidin 647 (detect F-actin). Super-resolution images were acquired using SMLM. (**A**) x-y cross section (600 nm thick in z) of an SMLM image of GFP-NiV-M (green) and F-actin (red). (**B** and **C**) Left column: the boxed regions in (**A**) are enlarged to show the detailed distribution of NiV-M and F-actin. Middle column: the cluster map of NiV-M (green). Right column: the corresponding pseudo-colored correlation map. The degree of correlation (DoC) value of individual NiV-M localizations with F-actin is shown in a heatmap, with −1 (segregation) in deep blue, and 1 (correlation) in deep red. Scale bars: 1 μm in (**A**); 200 nm in (**B** and **C**). (**D**) Correlation analysis. i) Degree of correlation of individual localizations from both channels. The density gradients of both channels are calculated along an increasing radius size (r) around each localization. The two density gradients are tested for correlation, resulting in a DoC value for each localization. The DoC value ranges from −1 to +1; with −1 indicating anti-correlation (segregation), 0 indicating random distribution, and +1 indicating correlation. ii) Localizations at the interface of the clusters (pink) have DoC values ≥ 0.4 (28). Therefore, the threshold of 0.4 identifies co-clustered localizations. iii) A NiV-M cluster was considered on the F-actin when the number of co-clustered localizations (DoC ≥ 0.4) exceeded 10 (26, 31). NiV-M clusters that did not meet these criteria were considered away from F-actin. (**E-G**) The diameter (**E**), relative density (**F**), and circularity (**G**) (The cluster is a true circle when circularity is 1) of NiV-M clusters on (n = 232) and off F-actin (n = 240) from 17 cells. Bars represent mean ± SD. *p* value was obtained using Welch’s *t*-test. ****: *p* ≤ 0.0001.

### F-actin retains NiV-M assembly sites at the plasma membrane

We envision two possible mechanisms by which F-actin promotes NiV-M nanoclustering: 1) F-actin facilitates the recruitment of new NiV-M molecules to the existing assembly sites; and 2) F-actin stabilizes existing assembly sites and prevents their disassembly or premature release. To distinguish between these two scenarios, we determined the assembly dynamics of GFP-NiV-M by monitoring the intensity changes of individual punctum over time using total internal reflection fluorescence (TIRF) microscopy combined with single-particle tracking (SPT) analysis.

NiV-M puncta started to appear on the PM at 6 hrs post-transfection in PK13 cells and accumulated to an optimal level for microscopic observation and SPT analysis between 12 and18 hrs post-transfection (Fig. 4A). The signals were observed either as diffused distributions or as discrete puncta (Fig. 4A), which aligns with our previous SMLM observations that NiV-M nanoclusters were small and uniform in size (perceived as diffused signals under conventional fluorescence microscopy) within the cell interior, whereas larger clusters were detected at the cell periphery (27). We observed a consistent increase in the intensity of diffused NiV-M and a slower growth of the number of NiV-M puncta (Fig. 4B), suggesting that NiV-M molecules were first expressed in the diffused form and then assembled into puncta. We selected cells that harbored a low number of puncta at the beginning of observation and monitored the dynamics of their fluorescence intensity change at the ventral plasma membrane over a 45-minute period at 5 s/frame using TIRF microscopy and SPT analysis. Since we observed membrane protrusions of ∼1 μm in length induced by NiV-M in COS-7 cells (Fig. 1E), we did not analyze the intensity of the elongated puncta because they were likely virus budding sites that were fully assembled or at a late stage of assembly. Further, we noticed that the NiV-M nanoclusters were well-spaced in SMLM (Fig. 3A) at the time of observation, and the puncta for intensity analysis were likely individual NiV-M assembly sites instead of large aggregations of NiV-M. Fig 4C is an averaged intensity profile of NiV-M puncta detectable over 75% of the total observation time. The intensity profile revealed three distinct phases: an initial rapid increase in fluorescence intensity (phase I), followed by a stationary phase (phase II) characterized by a stable intensity with minor fluctuations, and a final phase of fluorescence decay (phase III, Fig. 4C). To distinguish the intensity changes associated with the behaviors of assembly sites from the overall intensity alteration within the cells, we imaged 1) NiV-M, 2) an assembly-deficient mutant NiV-M-D339A (10), and 3) monomeric eGFP using epi-illumination to assess the overall intensity change. We observed a steady increase in the intensity of both diffused NiV-M wt and D339A, similar to the pattern seen with eGFP (Fig. 4D). By contrast, the intensity of NiV-M puncta follows a distinct three-stage pattern (Fig. 4C). Additionally, the intensity of diffused D339A increased more rapidly than that of the wt, suggesting that the diffused D339A was less efficient in aggregating into puncta, thereby confirming its assembly deficiency (Fig. 4D). These data inform us that 1) NiV-M puncta were derived from the assembly of diffused NiV-M at the PM (Fig. 4B and D), and 2) the rapid growth of fluorescence intensity of NiV-M puncta (phase I in Fig. 4C) was not due to an intensity change of the GFP signal (Fig. 4D), although phase III might be a combination of NiV-M puncta moving out of the TIRF illumination area (endocytosis or release) and photobleaching. Therefore, the NiV-M puncta intensity profile reflects three stages in the assembly and budding process: 1) the assembly stage where diffused NiV-M molecules are recruited to the PM and assemble into puncta; 2) the membrane-retention stage during which the NiV-M puncta remain on the PM with minimal changes in intensity; and 3) the release stage where the NiV-M puncta are either endocytosed or released into the extracellular space (Fig.4C).

**Fig. 4.**
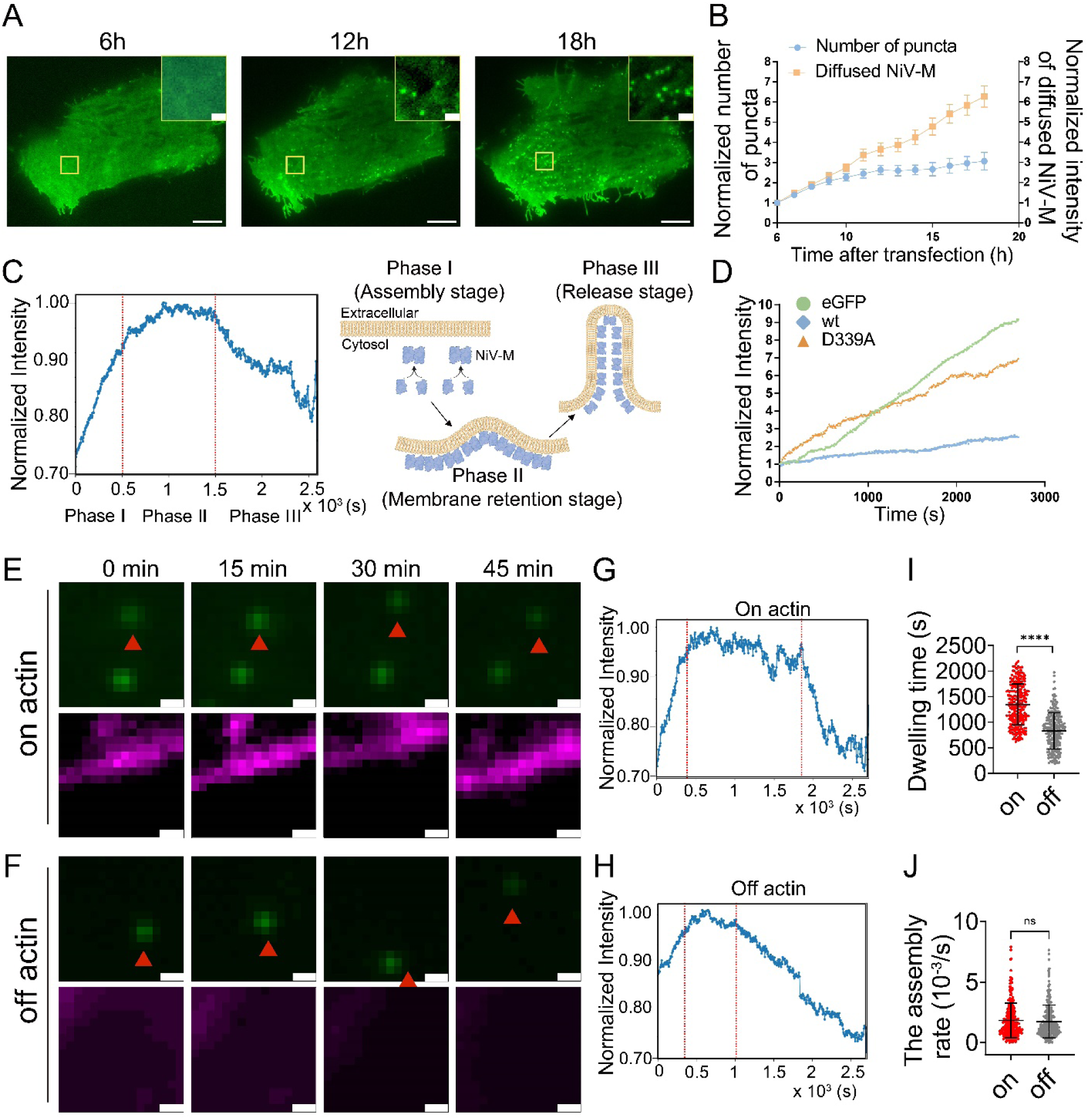
F-actin retains NiV-M assembly sites at the plasma membrane. (**A**) NiV-M puncta at the ventral plasma membrane PK13 cells expressing GFP-NiV-M was monitored by TIRF microscopy at 6 h, 12 h, and 18 h post-transfection. A representative image at each time point is shown. Scale bar: 10 μm and 1 μm. (**B**) PK13 cells expressing NiV-M were imaged 6-18 hrs post-transfection using epifluorescence illumination and TIRF. The number of puncta and intensity of diffused NiV-M are shown. (**C**) The intensity profile of GFP-NiV-M averaged from 250 tracks of 20 cells. Three phases were determined as shown in the “Materials and Methods” and separated by red dashed lines in the graph. (**D**) PK13 cells expressing monomeric eGFP, GFP-NiV-M-wt, or GFP-NiV-M-D339A were imaged over 45 minutes using epifluorescence illumination. The intensity change of GFP signals over time is shown. (**E, F**) Representative images of PK13 cells expressing both GFP-NiV-M-wt (green) and F-tractin-mCherry (magenta) at 0, 15, 30, and 45 minutes are shown. Scale bar: 1 μm. (**G** and **H**) The intensity change of individual NiV-M puncta was monitored by TIRF microscopy at 5 s per frame for 45 minutes and analyzed by SPT. Intensity profiles of “on-actin” (**G**) and “off-actin” (**H**) NiV-M tracks over 45 minutes. The total number of tracks is n = 260 (on-actin) and n = 238 (off-actin) from 30-40 cells per group. (**I**) The dwelling time of “on-actin” and “off-actin” NiV-M tracks. (**J**) The assembly rate of “on-actin” and “off-actin” NiV-M tracks. Bars represent mean ± SEM (B) mean ± SD (I and J). *p* value was obtained using Welch’s t-test. ns: *p* > 0.05, ****: *p* ≤ 0.0001.

By co-expressing Ftractin-mCherry, a probe for F-actin, and GFP-NiV-M in PK13 cells, we observed that some NiV-M puncta (“on actin”; Fig. 4E) stayed on or moved along F-actin, while some moved laterally away from F-actin (“off actin”; Fig. 4F). We classified the “on-actin” and “off-actin” tracks by assessing the percentage of events colocalized with F-tractin on individual tracks. A track was classified as “on-actin” if >98% of the events were colocalized with F-tractin. If <2% of the events were colocalized with F-tractin, the track was “off-actin”. The intensity profiles of “on-actin” (Fig. 4G) and “off-actin” (Fig. 4H) tracks were averaged from 200-300 tracks of each case. For the “on-actin” tracks, NiV-M puncta underwent distinctive assembly, membrane-retention, and release stages (Fig. 4G). However, the membrane-retention stage was short and not clearly distinguished from the release stage for the “off-actin” tracks (Fig. 4H). Our analysis of the duration of the membrane retention stage, measured by membrane dwelling time, shows that the “on-actin” tracks have a significantly longer membrane dwelling time than the “off-actin” tracks (Fig. 4I). Interestingly, the assembly rates, represented by the slope of the curve at the assembly stage, were similar between the “on-actin” and “off-actin” NiV-M tracks (Fig. 4J). These data indicate that F-actin stabilizes NiV-M assembly sites by preserving their association with the PM, rather than promoting the recruitment of additional NiV-M molecules to the assembly sites.

### The impaired F-actin interaction of NiV-M-I349A results in dispersed nanoclusters and reduced membrane retention time

To further investigate whether direct interaction with actin plays a role in NiV-M assembly, we tested the nano-organization of actin-binding-deficient NiV-M mutants on cell membranes. Single-color SMLM images show that I349A forms more dispersed and smaller clusters than that of the wt (Fig. 5A and 5B). We also observed a higher proportion of non-clustered I349A localizations compared to wt (Fig. 5A and 5B, cluster map), echoing the analysis showing a lower clustering tendency of I349A (Fig. 5C) and a lower percentage of the I349A localizations segregated into clusters (Fig. 5D). Similarly, clusters formed by I349A were smaller in size (Fig. 5E) but exhibited similar packing (Fig. 5F) and more regular shapes compared to the wt (Fig. 5G), although the total density was comparable to that of wt (Fig. 5H). These results suggest that disrupting NiV-M-actin interaction leads to a more dispersed distribution of NiV-M on the PM, highlighting a deficiency in the assembly process. By contrast, K351A did not significantly alter the nano-organization of NiV-M or result in any differences in the cluster morphology on the cell membrane (Fig. S3). Indeed, although K351A exhibited reduced VLP production compared to wt (Fig. 1D) and a noticeable decrease in actin association (Fig. 1C), we only observed a mild decrease in membrane protrusion (Fig. 1E). We speculate that the K351A mutation may contribute to fission of the budding VLPs. We also analyzed the intensity change of the puncta formed by I349A using TIRF and SPT, and compared it to that of wt as described above (Fig. 5I-K). Interestingly, while the assembly rate of I349A was comparable to that of wt (Fig. 5M), its membrane dwelling time was markedly reduced (Fig. 5L). These data show that the I349 residue, critical for NiV-M-actin interaction, maintains NiV-M nanoclustering and membrane association, further supporting the F-actin-dependent integrity of NiV assembly sites at the PM.

**Fig. 5.**
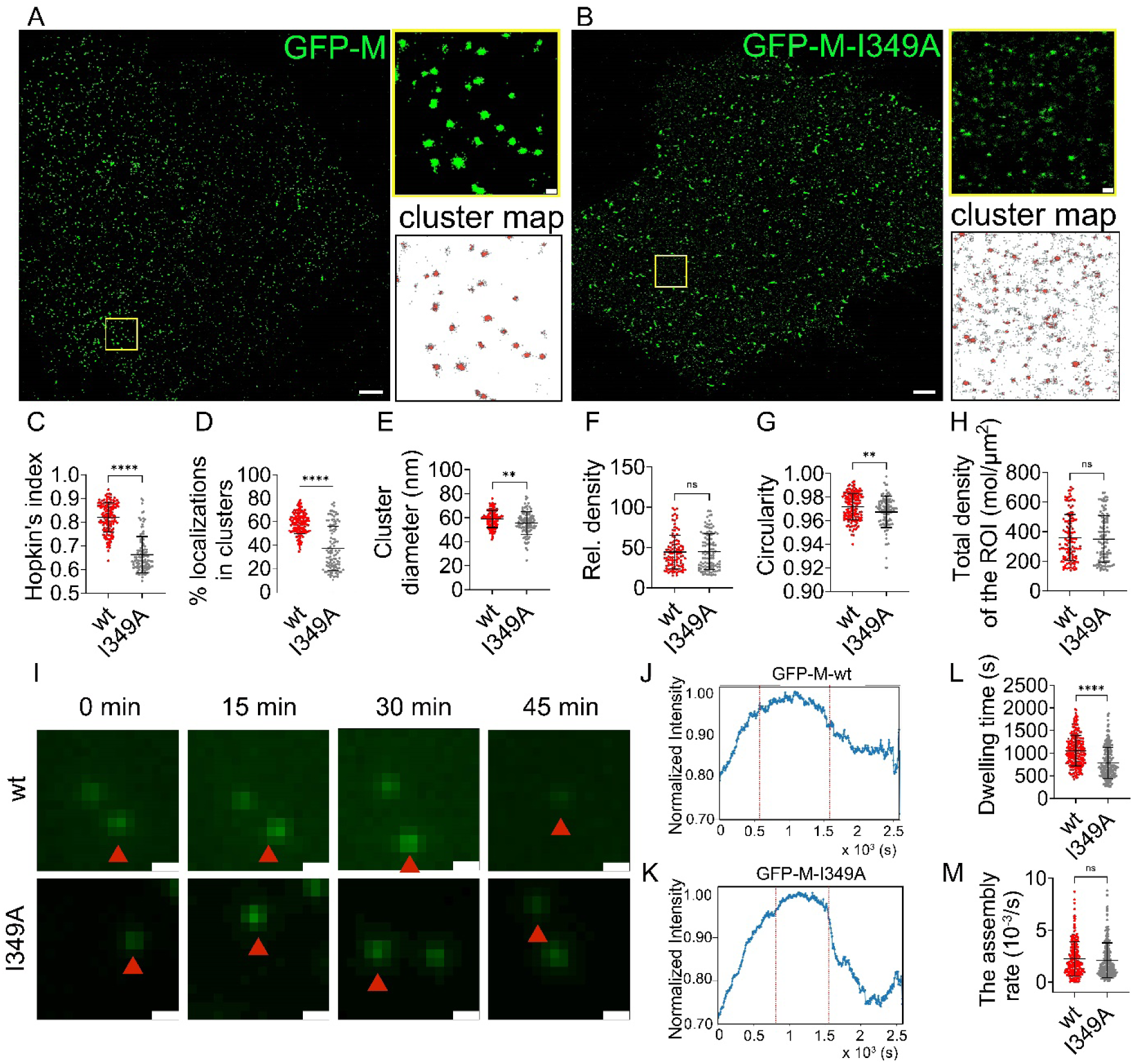
Mutations at the actin-binding domain in NiV-M alters the nano-organization of NiV-M and assembly dynamics. PK13 cells were transfected with GFP-tagged NiV-M-wt or I349A. NiV-M constructs were stained using a goat anti-GFP antibody and donkey anti-goat antibody conjugated to Alexa Fluor 647. (**A** and **B**) x-y cross-section (600 nm thick in z) of the SMLM images of GFP-tagged NiV-M-wt (**A**) and I349A (**B**) at the ventral membrane of PK13 cells. The boxed regions are enlarged, and the cluster maps of the NiV-M localizations are shown. Scale bars: 1 μm and 200 nm. (**C**) The Hopkin’s index of the localizations of NiV-M-wt and I349A. (**D-H**) The percentage of localizations in clusters (**D**), cluster diameter (**E**), relative density (**F**), circularity (**G**), and total density of the ROI (**H**). Sample size n = 150 (wt) and 106 (I349A) from 12-18 cells per group. (**I**) Representative images of PK13 cells expressing GFP-tagged NiV-M-wt and I349A at 0, 15, 30, and 45 minutes. Scale bar: 1 μm. (**J** and **K**) The intensity change of individual NiV-M-wt or I349A puncta was monitored by TIRF microscopy at 5 s per frame for 45 minutes and analyzed by SPT. Intensity profiles of NiV-M-wt (**J**) and I349A (**K**) over 45 minutes. The total tracks are n = 232 (wt) and n = 254 (I349A) from 30-40 cells per group. (**L**) The dwelling time of NiV-M-wt and I349A tracks. (**M**) The assembly rate of NiV-M-wt and I349A tracks. Bars represent mean ± SD. *p* value was obtained using Welch’s t-test. ns: *p* > 0.05, **: *p* ≤ 0.01, ****: *p* ≤ 0.0001.

### Arp2/3-driven F-actin branching retains NiV-M assembly sites on the membrane and promotes VLP budding and release

Our findings indicated that F-actin primarily functions to stabilize NiV-M assembly sites at the PM after assembly is complete, rather than facilitating the recruitment of additional NiV-M molecules to these sites. Since Arp2/3 complex is known to drive F-actin branching and is important for both virus entry and escape (14), we hypothesized that Arp2/3-driven F-actin branching and polymerization are responsible for the membrane retention of NiV-M assembly sites. We observed that the number of membrane protrusions in COS-7 cells stably expressing NiV-M was inversely correlated with the treatment dosage of an Arp2/3-inhibitor, CK-666 (Fig. 6A). We also observed many finger-like structures formed by NiV-M in these COS-7 cells by TIRF at 6-18 hrs post-transfection (Fig. 6B). These finger-like structures closely resembled the membrane protrusions seen in SEM images (Fig. 6A). The ratio of the number of finger-like structures to that of puncta increased over time in the control group (DMSO), suggesting that the finger-like structures likely developed through the transition of puncta (Fig. 6C). By contrast, we did not observe an increase of the finger-like structures over puncta in COS-7 cells treated by CK-666 (Fig. 6C). This observation suggests that NiV-M hijacks the Arp2/3-mediated actin debranching for budding. Indeed, a dose-dependent decrease in VLP production in HeLa cells treated with CK-666 was observed (Fig. 6D). In comparison, VLP production remained at similar levels in cells treated with a CK-666 inactive analog, CK-689 (Fig. 6D). Furthermore, siRNA-mediated knockdown of the endogenous Arp2 and Arp3 decreased VLP production in HeLa cells stably expressing NiV-M. These results suggest that NiV-M co-opts the Arp2/3 complex for budding and VLP production (Fig. 6E). To further investigate whether the Arp2/3 complex affected the membrane retention of NiV-M assembly sites, we analyzed the NiV-M intensity profile using TIRF and SPT in PK13 cells treated with 200 μM CK-666. This treatment resulted in a decrease in VLP production without substantial cytotoxicity effects in PK13 cells (Fig. S4). Interestingly, a profound reduction in the membrane-dwelling time of NiV-M puncta was observed in cells treated with CK-666 compared to control-treated cells (Fig. 6F and G), suggesting that the Arp2/3-driven actin debranching is critical for membrane retention of NiV-M puncta. Additionally, it is unlikely that Arp2/3 is substantially involved in the assembly stage since the assembly rate was not affected by CK-666 treatment (Fig. 6F and H). Taken together, these results highlight the role of the Arp2/3 complex in promoting NiV VLP production by driving actin polymerization and branching during budding. This process is facilitated by F-actin-dependent membrane retention, which may result from the interaction between NiV-M and the Arp2/3 complex.

**Fig. 6.**
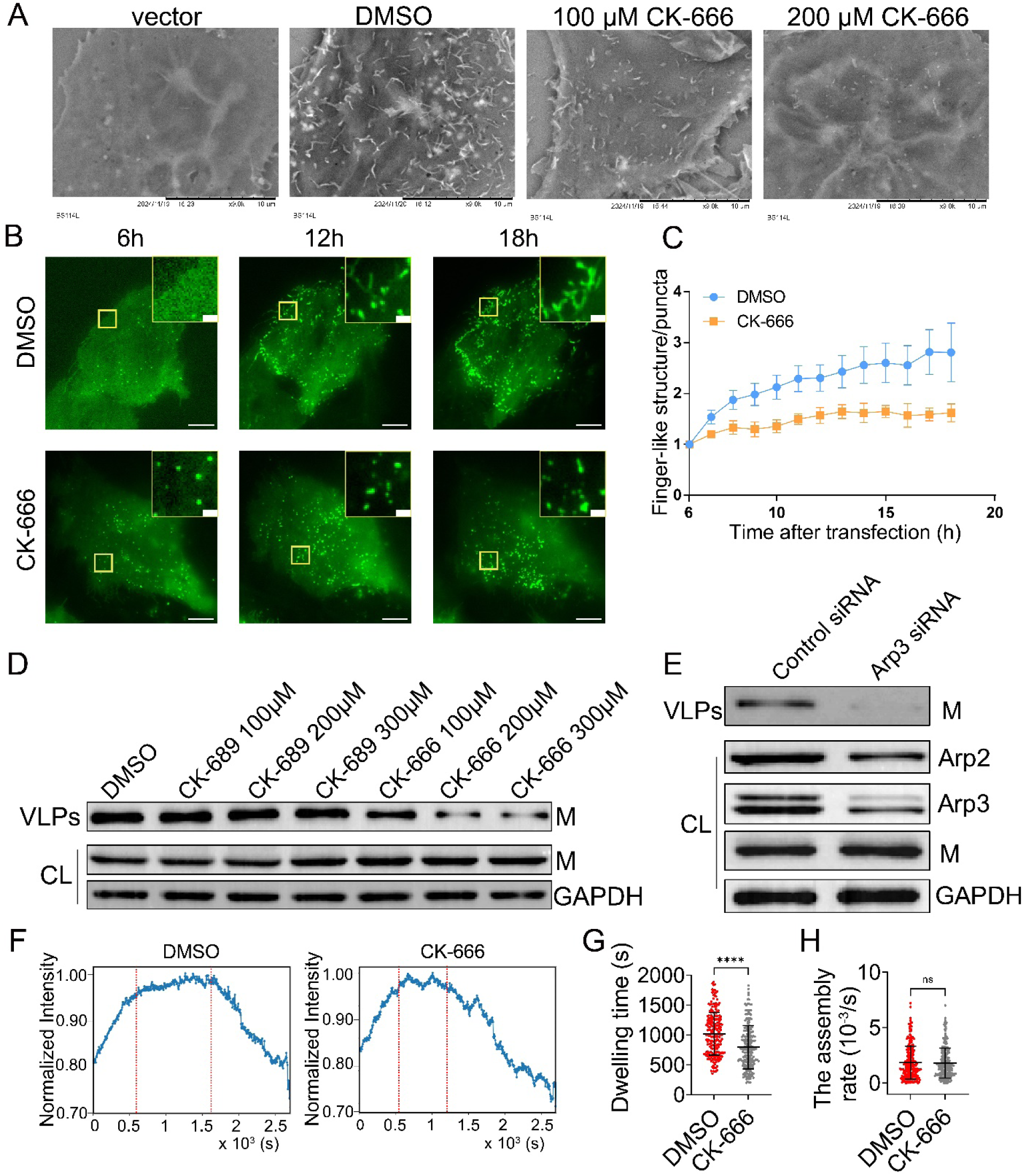
Arp2/3-driven F-actin branching retains NiV-M assembly sites on the membrane and promotes VLP budding and release. (**A**) COS-7 cells stably expressing 3×FLAG-NiV-M treated by DMSO, 100 μM, and 200 μM CK-666 were imaged by SEM. COS-7 cells transduced with empty pQCXIP vector were used as a control. Representative images from at least three independent experiments are shown. Scale bar: 10 μm. (**B**) PK13 cells expressing GFP-NiV-M were treated with DMSO and 200 μM CK-666 at 6 hrs post-transfection. The morphology of NiV-M assembly sites at the ventral plasma membrane was monitored by TIRF microscopy at 6 h, 12 h, and 18 h post-transfection. A representative image at each time point is shown. Scale bar: 10 μm and 1 μm. (**C**) The ratio of the number of finger-like structures to puncta was determined and tracked at 6-18 hrs post-transfection in PK13 cells treated with either DMSO or CK-666. (**D**) Western blot analysis of VLPs and cell lysates from HeLa cells stably expressing 3×FLAG-NiV-M, treated by CK-689 and CK-666. GAPDH is used as a loading control. NiV-M was detected using a mouse anti-FLAG antibody and GAPDH mouse anti-GAPDH antibody. Proteins were detected by HRP-conjugated secondary antibodies. Results from at least three independent experiments were shown. (**E**) HeLa cells stably expressing 3×FLAG-NiV-M were transfected by a control siRNA and an Arp3-targeting siRNA. Western blot analysis was performed on cell lysate and VLPs at 48 hrs post-transfection. NiV-M was detected using a mouse anti-FLAG antibody, Arp2 a rabbit anti-Arp2 polyclonal antibody, Arp3 a rabbit anti Arp3 monoclonal antibody, and GAPDH mouse anti-GAPDH antibody. Proteins were detected by HRP-conjugated secondary antibodies. Results from at least three independent experiments were shown. (**F**) PK13 cells were transfected with GFP-NiV-M and treated by 200 μM DMSO or CK-666 drug. The intensity change of individual NiV-M assembly sites was monitored by TIRF microscopy at 5 s per frame for 45 minutes and analyzed by SPT. The intensity profiles of NiV-M in DMSO or CK-666-treated PK13 cells were plotted. Sample size n = 287 (DMSO) and 277 (CK-666) tracks from 30-40 cells per group. (**G**) The dwelling time of NiV-M puncta on plasma membrane in DMSO- and CK-666 treated cells. (**H**) The assembly rate of NiV-M puncta in DMSO- and CK-666 treated cells. Bars represent mean ± SEM (C) mean ± SD (G and H). *p* value was obtained using Welch’s t-test. ns: *p* > 0.05, ****: *p* ≤ 0.0001.

## Discussion

In this study, we employed a combination of SMLM, live cell imaging and SPT, electron microscopy, and biochemistry analysis to dissect the processes of NiV assembly and budding. Our findings reveal that NiV-M utilizes F-actin to sustain the nanoscale organization and membrane association of assembly sites at the PM. Notably, we show that Arp2/3-driven F-actin branching and polymerization play a critical role in maintaining NiV-M assembly sites at the membrane. Furthermore, NiV-M co-opts Arp2/3-driven actin branching to facilitate the formation of membrane protrusions during the budding process.

NiV-M forms nanoclusters that are stabilized by host F-actin at the PM. SMLM images show that NiV-M forms nanoclusters at the membranes of host cells and VLPs (Fig. 2) (27, 29). This is supported by several recent cryo-electron microscopy studies showing that paramyxovirus M proteins are organized into a two-dimensional paracrystalline array associated with the PM of infected cells and cell-free viral membranes (7, 8). However, only 10% of NDV virions contain a visible layer formed by the M protein, and this layer is often discontinuous in many virions (7). These observations suggest that 1) both ordered and disordered M protein arrays can be incorporated into viruses and 2) the paracrystalline array may undergo a structural rearrangement during budding. This rearrangement may result from the different composition and organization of host factors in host cells and virus particles. Actin has been identified as an interacting partner of NiV-M and found in NiV VLPs produced by transfection of NiV-F, G, and M, highlighting the critical role of actin in the assembly and budding of viruses (24, 32, 33). Our data revealed an interaction between actin and a C-terminal motif in NiV-M, similar to that in SeV-M (18). Disruption of this interaction through point mutations at I349 and K351 reduced VLP production (Fig. 1), highlighting the functional importance of this motif in the viral assembly process. SMLM imaging shows that close proximity to or direct interaction with F-actin enhances NiV-M clustering, resulting in larger and more irregularly shaped clusters that resemble viral assembly sites (Fig. 3 and 5). On the contrary, a recent TIRF/dSTORM study shows that HIV-1 Gag assembly sites prefer a low local F-actin density resulting from actin branching inhibition (23). This discrepancy may result from different experiment systems, but more importantly, it suggests diverse roles of F-actin in virus assembly and budding. F-actin may form an actin cortex underneath the PM as a barrier to virus release, or be actively recruited by viral proteins to generate membrane protrusions that facilitate virus budding.

Virus assembly, budding, and release are closely related processes, especially for viruses that assemble and bud at the PM, such as paramyxoviruses, retroviruses, and orthomyxoviruses (34). We distinguished these processes by tracking individual NiV-M assembly sites. The intensity profile of NiV-M assembly sites was divided into three phases that represent assembly, membrane retention, and release stages, respectively (Fig. 4C). Our data show that the dissociation of NiV-M puncta from F-actin (Fig. 4), the disruption of NiV-M-F-actin interaction by the I349A mutation (Fig. 5), and the inhibition of the Arp2/3-driven actin branching (Fig. 6) resulted in a shorter membrane-retention stage. Together with the results showing that Arp2/3 promoted VLP production by facilitating the generation of NiV-M induced membrane protrusions (Fig. 6), our data demonstrate that 1) NiV-M assembly sites remain on the PM after assembly completion (Fig.4); 2) F-actin is important in this retention by preventing NiV-M disassembly (Fig. 2-4); 3) membrane retention of NiV-M assembly sites is likely linked to F-actin polymerization and branching by Arp2/3 (Fig. 6); and 4) membrane retention, in turn, promotes the NiV-M-F-actin-Arp2/3 interactions for membrane protrusion generation for virus budding. We speculate that the membrane-retention stage is crucial for the successful budding of virions because NiV-M recruits essential host factors for budding during this stage, such as F-actin and the Arp2/3 complex. Defects at this stage will lead to the NiV-M assembly sites being endocytosed and/or prematurely released. A previous study also tried to pinpoint the mechanisms underlying individual phases in HIV-Gag assembly. HIV virions lacking Vpu, a protein essential for counteracting tetherin-mediated restriction of virus release, were used to characterize the phase associated with viral release. However, no significant differences in assembly rate or the duration of each phase were observed between wt and Vpu-deficient virions. This was largely because tetherin-mediated restriction occurs primarily after the completion of the budding events (35).

A recent study shows that NiV-M associates with PS and PI(4,5)P_2_ on the PM, and this association induces conformational changes in the surface of the M dimer that trigger membrane deformation (10). We show that NiV-M requires the Arp2/3 complex-induced F-actin remodeling to generate membrane protrusion (Fig. 6). These data imply that membrane deformation caused by conformational changes of NiV-M may trigger F-actin remodeling by the Arp2/3 complex. Membrane deformation is known to cause actin polymerization and the accumulation of signaling proteins involved in actin remodeling. For example, local accumulation of F-actin was observed on various vertical nanostructures and the location of this accumulation is curvature-dependent (36). Interestingly, the actin regulator WASP enriches at sites of beads-induced membrane deformation and facilitates the recruitment of the Arp2/3 complex to these sites, stimulating actin assembly that couples membrane deformation to cytoskeleton remodeling (37). It is likely that NiV-M induces budding by combining membrane deformation—caused by its binding to PS and PI(4,5)P_2_—with Arp2/3-driven F-actin assembly.

Remodeling of cortical actin during virus assembly and budding has been reported for many viruses, although its precise effects remain a topic of ongoing debate. While the majority of evidence comes from endpoint biochemistry analysis, including the observation of incorporation of actin regulators in newly formed virions, the assessment of virus release upon knockdown of key actin regulators, and the disruption of cytoskeleton dynamics using drug treatment (38), recent advances in super-resolution imaging and dynamic analysis have provided structural and temporal details of the underlying mechanisms (23, 39, 40). Notably, a recent study on HIV-infected T cells shows that HIV Gag prefers to assemble at areas with low F-actin density at the PM and decreased actin branching induced by Arp2/3 favors HIV Gag assembly and virus production (23). However, the exact step in assembly affected by F-actin density and Arp2/3 remains unclear. By combining SMLM and live cell imaging, our data show that the Arp2/3 complex promotes NiV VLP release by facilitating F-actin assembly and branching, resulting in the generation of membrane protrusions. Overall, these findings highlight a diverse role of actin remodeling in virus assembly and budding, which can be context-dependent and influenced by specific viral and host factors. Future development of minimal in vitro systems, such as non-infectious viral replicons, will be valuable for validating these findings on NiV, particularly given the stringent high-containment requirements for studying this virus.

## Materials and Methods

### Plasmids

The plasmids expressing monomeric enhanced GFP (GFP)- and 3xFLAG-tagged NiV-M were adapted and modified from a previous study (41). NiV-M, NiV-M-I349A, and NiV-M-K351A were produced by site-directed mutagenesis using pcDNA 3 plasmids coding for GFP- and FLAG-tagged NiV-M. The plasmid expressing pLV-Ftractin-mCherry (Addgene plasmid # 85131) is a gift from Tobias Meyer. The Ftractin-mCherry gene was ligated into a pcDNA3 expression plasmid flanked by KpnI and EcoRI restriction sites. For stable cell line establishment, cDNA fragments coding for FLAG-tagged NiV-M, NiV-M-I349A, and FLAG-NiV-M-K351A gene were ligated into a pQCXIP expression plasmid flanked by PacI and EcoRI restriction sites, respectively. MLV-Gag/Pol was a gift from Dr. Chen Liang at McGill University. The VSV-G envelope expressing plasmid, pmd2.G, was a gift from Didier Trono (Addgene plasmid # 12259).

### Cell lines and tissue culture

PK13, COS-7, HEK293T, and HeLa cells were cultured at 37°C and 5% CO_2_ in DMEM (Sigma-Aldrich, D6429) complemented with 10% fetal bovine serum (Invitrogen, 12483-020). Cells were passaged using phosphate-buffered saline (PBS, Invitrogen, 10010-049) and 0.25% Trypsin-EDTA solution (Invitrogen, 25002-072). Cells were monitored routinely for mycoplasma contamination using a mycoplasma detection PCR kit (ABM, G238). To generate stable cell lines, retroviral pQCXIP vectors encoding FLAG-tagged NiV-M constructs (4 μg), MLV-Gag/Pol (4 μg), and pmd2.G (0.5 μg) plasmids were co-transfected using polyethylenimine (PEI) at 1 mg/mL (Polysciences, 23966-100) to HEK293T cells cultured in a 10 cm dish. Viral supernatant was collected 72 hrs post-transfection to infect COS-7 or HeLa cells. After 3 days of infection, puromycin selection was applied for 21 days to establish a stable cell line expressing the FLAG-tagged NiV-M constructs.

### Drugs, siRNA, and antibodies

Drugs used in this study are: Latrunculin A (Calbiochem, 428021), CK-666 (Calbiochem, 182515), CK-689 (Calbiochem, 182517) and DMSO (Sigma-Aldrich D8418-100ML). Antibodies used in this study are: anti-FLAG mouse monoclonal antibody (Sigma-Aldrich, F1804), anti-GFP goat antibody (Abcam, ab5450), anti-GFP rabbit antibody (Invitrogen, G10362), anti-β-actin mouse antibody (Sigma-Aldrich, A2228), anti-GAPDH mouse antibody (Sigma-Aldrich, CB1001), anti-Arp3 rabbit antibody (Abcam, ab181164), anti-Arp2 rabbit antibody (Abcam, ab129018), anti-Vimentin rabbit antibody (Abcam, ab92547), anti-goat donkey polyclonal antibody, Alexa Fluor^®^ 647 conjugated (A21447), anti-rabbit donkey polyclonal antibody, Alexa Fluor^®^ 647 conjugated (A16025), phalloidin Alexa Fluor 647 (A22287), anti-goat donkey polyclonal antibody, HRP conjugated (Jackson Immunoresearch, 705-035-147); Anti-mouse goat polyclonal antibody, HRP conjugated (Biorad, 1705047); Anti-rabbit goat polyclonal antibody, HRP conjugated (Biorad, 1706515). The cy3B (Cytiva, PA63101) is conjugated to the donkey anti-goat antibody (Jackson ImmunoResearch, 705-005-003) and donkey anti-rabbit antibody (Jackson ImmunoResearch, 711-005-152) in-house by Ablabs. The siRNAs used in this study are: siGENOME SMARTpool human ACTR3 (M-012077-01-0005) and non-targeting siRNA pool (D-001206-13-05).

### Virus-like-particle (VLP) production and purification

To determine the VLP production of various NiV-M constructs, HEK293T cells were transfected by FLAG-tagged- or GFP-tagged NiV-M constructs and pcDNA3 vector at a 1:2 ratio by using PEI at 1 mg/ml. At 48 hrs post-transfection, the supernatant of the cell culture was collected for VLP purification. To study the effect of Arp2/3 on NiV VLP production, PK13 or HeLa cells stably expressing FLAG-tagged NiV-M were seeded in 6-well plates, and treated with DMSO, 100, 200, and 300 μM of CK-666 and CK-689 at 6 hrs after seeding. After 48 hrs of drug treatment, the supernatant was collected for VLP purification. Alternatively, HeLa cells stably expressing FLAG-tagged NiV-M were transfected by 20 nM anti Arp2/3 siRNA using 3 μl RNAiMAX (Invitrogen, 13778075). At 48 hrs post-transfection, the supernatant of the cell culture was collected. For VLP purification, the supernatant was filtered using 0.45 μm PES membrane (VWR, 76479-020) and subjected to ultracentrifugation on a 20% sucrose cushion at 125, 392 rcf with an SW 41 rotor (Beckman) for 90 minutes. The VLP-containing pellets were resuspended in 5% sucrose-NTE buffer and stored in −80°C. The cells from each different treatment were lysed, and both cell lysates and purified VLPs were subjected to Western blot analysis to evaluate VLP production activity.

### Western blot analysis

The virus-producing cells were lysed using RIPA buffer (Sigma-Aldrich, 20-188) supplemented with EDTA-free protease inhibitor cocktail (Sigma-Aldrich, 11836170001). The cell lysates were collected after centrifuge at 16,000 ×g for 20 minutes at 4°C. VLP and cell lysate were supplemented with 1x SDS loading dye [60 mM Tris-HCl (pH=6.8) (Tris base, Millipore, 648311-1kg); 2% SDS (Sigma-Aldrich, L3771); 10% glycerol (Sigma-Aldrich, G5516-1L), 0.025% Brophenol blue (Sigma-Aldrich, 114391-5G)] and 15 mM DTT (Thermo Scientific, R0861), and heated at 95°C for denature. The denatured VLP and cell lysates were separated on a 10% polyacrylamide gel for SDS-PAGE. Proteins were then transferred to an activated polyvinylidene difluoride (PVDF) membrane (Millipore, IPVH00005), pore size 0.45 μm. The PVDF membrane was then blotted using 1% Bovine Serum Albumin (BSA, Sigma-Aldrich, A9647) blocking buffer in PBS for 90 minutes at room temperature and incubated with primary antibodies. An HRP-conjugated goat anti-mouse, HRP-conjugated goat anti-rabbit, or donkey anti-goat secondary antibody was used for protein detection. The membranes were then developed with Biorad Western ECL substrate (Bio-rad, 170560). Membrane images were acquired using Chemic Doc MP Imaging System (BioRad). The integrated intensity of the NiV-M bands was measured by densitometry using ImageJ. The budding index is the ratio of the normalized amount of NiV-M in the VLPs over that of the cell lysates. The ratio of the NiV-M group is set as 100% (10).

### Co-immunoprecipitation

HEK293T cells were transfected with the following plasmids: 1) 3xFLAG-NiV-M-wt, 2) 3xFLAG-NiV-M-I349A, and 3) 3xFLAG-NiV-M-K351A. At 48 hrs post-transfection, cells from 1 well of a 6-well plate were washed with PBS and lysed in 200 μl lysis buffer provided with the μMACS DYKDDDDK isolation kit (Miltenyi Biotec, 130-101-591), and supplemented with protease inhibitors. Cells were isolated on ice for 30 minutes. Cell debris was removed by centrifuge at 16,000 x g for 20 minutes at 4°C. 60 μl of cell lysate was set aside for immunoblot analysis, and the rest was used for immunoprecipitation, as recommended by the manufacturer. 6 μl anti-DYKDDDDK microbeads (Miltenyi Biotec, 130-101-591) were added to 140 μl cell lysates and incubated for 30 minutes on ice. μ columns (Miltenyi Biotec, 130-042-701) were prepared according to the manufacturer’s instructions. Lysate was run over the columns, and microbeads were washed according to the manufacturer’s instructions. 20 μl preheated elution buffer (95°C) was added to the column before eluting the bound immunoprecipitated protein in 50 μl elution buffer. Elute was separated by 10% SDS-PAGE, and proteins were immunoblotted by mouse anti-FLAG and mouse anti-β-actin antibodies. The HRP-conjugated goat anti-mouse and goat anti-rabbit secondary antibodies were used for protein detection. The integrated intensity of the NiV-M and β-actin band was measured by densitometry using ImageJ. The % actin pulldown is the ratio of β-actin over NiV-M in the IP group over that of the Input group. The ratio of the NiV-M group is set as 100%.

### Immunofluorescence for confocal microscopy

To visualize protein composition in VLPs, VLPs expressing GFP-NiV-M were bound to coverslips (Marienfeld #1.5H, 18 mm) coated by 2.5 μg fibronectin at 37°C for 4 hrs, followed by fixation using PBS containing 4% PFA. VLPs were incubated with BlockAid blocking solution (Life Technologies, B10710) at room temperature for 1 hr, and stained with mouse anti-β-actin and donkey anti-mouse antibody conjugated to Alexa Flour 647®. The percentage of VLPs containing β-actin was determined as a ratio of the number of GFP+/ β-actin+ particles over the number of GFP+ particles. Images were acquired using a laser scanning confocal microscopy Zeiss LSM710 and data analysis was performed using Imaris (Oxford Instruments).

### Immunofluorescence (IF) for SMLM

For SMLM imaging, 1×10^5^ PK13 cells were seeded on coverslips (Marienfeld #1.5H, 18 mm) coated with 2.5 μg fibronectin (Sigma-Aldrich, F4759-2 mg) in a 12-well plate, and transfected with 0.3 μg GFP-NiV-M constructs with 0.7 μg pcDNA 3 plasmids using lipofectamine 3000 (Invitrogen, L3000015) on the following day. In the case of drug treatment, PK13 cells expressing GFP-NiV-M were treated by using 1 µM DMSO or Latrunculin A (LatA) for 5 minutes before fixation. The protocol for immunofluorescence of actin cytoskeleton was adapted from Huang et al. (42).

For dual-color imaging of actin cytoskeleton and NiV-M constructs, cells were fixed at 20-24 hrs post-transfection with 0.3% glutaraldehyde (Sigma-Aldrich, G5882-50 ml) and 0.25% Triton X-100 (Sigma-Aldrich, T8787-50ML) in cytoskeleton buffer [CB buffer; 10 mM MES (Sigma-Aldrich, 475893) pH 6.1, 150 mM NaCl (Sigma-Aldrich, S9888-1KG), 5 mM EGTA (Sigma-Aldrich, E3889-10G), 5 mM glucose (Sigma-Aldrich, G8270-1KG), and 5 mM MgCl_2_ (Sigma-Aldrich, M8266-100G)] for 1-2 minutes, followed by a second fixation step using 400 µl of 2% glutaraldehyde (Sigma-Aldrich, G5882-50 ml) in CB for 10 minutes. Cells were treated with 1 ml 0.1% NaBH4 (Sigma-Aldrich, 452882-5G) (freshly prepared in PBS) for 7 minutes to reduce background fluorescence. For immunofluorescence of NiV-M, cells were fixed with PBS containing 4% paraformaldehyde (PFA; Electron Microscopy Sciences; 50980487) and 0.2% glutaraldehyde for 90 minutes at room temperature followed by permeabilization using 0.1% Triton X-100 in PBS. After fixation and permeabilization, cells were incubated with signal enhancer image-IT-Fx (Life Technologies, I36933) for 30 minutes at room temperature, and then blocked using BlockAid blocking solution for 1 hr at room temperature. The GFP-tagged NiV-M and mutants were detected by an anti-GFP rabbit (Invitrogen, G10362) or goat antibody (Abcam, ab5450) and a cy3B-conjugated donkey anti-rabbit or anti-goat secondary antibody. Cells were incubated with primary antibody overnight at 4°C, and then with the secondary antibody for 1 hr at room temperature. Each antibody incubation was followed by five PBS washes, 5 minutes each time. The F-actin was stained using 0.5 µM of Alexa 647-phalloidin (Invitrogen A22287), followed by three PBS washes. Cells were then fixed in PBS containing 4% PFA for 10 minutes at room temperature.

### SMLM setup, imaging, and data analysis

SMLM was performed on a custom-built microscopy described previously (26, 29). Briefly, the microscopy was built upon an apochromatic TIRF oil-immersion objective lens (Nikon, 60x; numerical aperture 1.49). Four lasers were used for excitation: a 639 nm laser (MRL-FN-639, 500 mW) for exciting Alexa Fluor^®^ 647, a 532 nm laser (MGL-III-532-300 mW) for exciting cy3B, a 488 nm laser (MBL-F-473-300 mW) for exciting GFP, and a 405 nm laser (MDL-III-405-100 mW) for reactivating Alexa Fluor 647 and cy3B. The emission fluorescence was separated using appropriate dichroic mirrors and filters (Semrock) and detected by electron-multiplying charge-coupled devices (EMCCD; Ixon, Andor). A feedback loop was employed to control the sample drift to <1 nm laterally and 2.5 nm axially. Fluorescence beads (Life technologies, F8799) were added to samples as fiducial markers for drift control. Samples were immersed in imaging buffer [TN buffer (50 mM Tris (pH 8.0) and 10 mM NaCl), 0.5 mg/ml glucose oxidase (Sigma-Aldrich, G2133-50KU), 40 μg/ml catalase (Sigma-Aldrich, C100), 10% glucose, and 50 mM mercaptoethylamine (MEA) (Sigma-Aldrich, 641022) or 143mM 2-Mercaptoethanol (Sigma-Aldrich, 444203) for dual-color SMLM imaging]. For SMLM imaging, samples were exposed to a laser power density of 1 kW/cm^2^ for the 639 nm and 532 nm lasers to activate Alexa Fluor® 647 and cy3B, respectively. A total of 40,000 images were acquired at 50 Hz to reconstruct one SMLM image. Custom-written software in MATLAB (Mathworks) was used to reconstruct SMLM images. Clusters of NiV-M localizations were identified and characterized using ClusDoC (31). The minute points and ε for DBSCAN were set at 4 and 20, respectively(26). The degree of colocalization assay (DoC) was performed using ClusDoc. A DoC value is generated for individual localizations. The colocalized clusters contain more than 10 localizations with a DoC value greater than 0.4 (28).

### Cytoskeleton extraction assay

We used a protocol described previously to extract the cytoskeleton and determine the association of NiV-M on the cytoskeleton (43). PK13 cells were transfected with FLAG-tagged-NiV-M for 48 hrs and subsequently washed with PBS. The cells were then incubated with cellular extraction buffer [50 mM PIPES (Sigma-Aldrich, P1851-25G), 50 mM NaCl, 5% Glycerol, 0.1% NP-40 (Sigma-Aldrich, NP40S-100ML), 0.1% Triton X-100 and 0.1% Tween 20 (Sigma-Aldrich, P9416-100ML)] and cytoskeleton wash buffer (50 mM Tris-HCl, pH 7.5) to obtain the soluble compartment (S). For the nuclear compartment (N), cells were incubated with nuclear extraction buffer [10 U/mL Benzoase nuclease (Sigma-Aldrich, E1014-5KU), 10 mM MgCl_2_ and 2 mM CaCl_2_ (Sigma-Aldrich, 21115-1ML) in 50 mM Tris-HCl buffer, pH 7.5]. Next, cells were washed with cellular extraction buffer and incubated with cytoskeletal solubilization buffer (1% SDS) to isolate the cytoskeletal compartment (C). The association of 3xFLAG-NiV-M with N, C, and S compartments was determined by Western blot analysis. Vimentin served as a control for compartment C, and GAPDH served as a control for compartments S and N. The extracts were supplemented with 1x SDS loading dye and 15 mM DTT, and heated at 95°C for denature. Then the extracts were separated on a 10% polyacrylamide gel for SDS-PAGE. Proteins were then transferred to an activated PVDF, pore size 0.45 μm. The PVDF membrane was then blotted using 1% Bovine Serum Albumin blocking buffer in PBS for 90 minutes at room temperature and incubated with primary antibodies (mouse anti-FLAG antibody, rabbit anti-Vimentin antibody, and mouse anti-GAPDH antibody). An HRP-conjugated goat anti-mouse or HRP-conjugated goat anti-rabbit was used for protein detection. The membranes were then developed with Biorad Western ECL substrate. Membrane images were acquired using Chemic Doc MP Imaging System.

### Scanning electron microscopy

COS-7 cells stably expressing FLAG-tagged NiV-M wt or mutants were seeded on coverslips (Marienfeld #1.5H, 12 mm) coated with 1.25 μg fibronectin (Sigma-Aldrich, F4759-2 mg) in a 24-well plate. After 24 hrs, cells were fixed with 2.5% (w/v) glutaraldehyde in 0.1 M sodium cacodylate buffer (Sigma-Aldrich, 70114-10ML-F) for 30-60 minutes at room temperature, followed by three PBS washes, 10 minutes each time. Then cells were fixed in pre-cooled 1% (v/v) osmium tetroxide (Sigma-Aldrich, 251755-5ML) for 30 minutes at room temperature, followed by three PBS washes. Cells were then dehydrated through a graded ethanol series (25%, 50%, 75%, 95%, and 100%) and dried using a Tousimis 931 critical point dryer (Leica EM CPD300). Sputter coating was conducted using a sputter coater (Leica EM ACE200). Images were captured at 9000 × magnification using scanning electron microscopy (Hitachi TM-1000).

### TIRF microscopy (TIRF) and single-particle tracking (SPT)

For SPT experiments, PK13 cells transfected by GFP-tagged NiV-M constructs or GFP-NiV-M and Ftractin-mCherry were subjected to TIRF-M imaging at 12-18 hrs post-transfection. In case of CK-666 treatment, 200 uM CK-666 was added to the growth medium of PK13 cells transfected by GFP-NiV-M at 6 hrs post-transfection and subjected to TIRF imaging at 12-18 hrs post-transfection. Imaging was performed by a Zeiss TIRF microscopy equipped with a 63x NA1.46 objective lens at TIRF illumination. A live cell chamber (Tokaihit) maintained a 37°C and 5% CO_2_ atmosphere. Images were acquired at 5 s per frame for 45 minutes. To analyze the fluorescence intensity change of individual NiV assembly sites, we used SPT analysis to track NiV-M puncta over time. The image analysis was performed in 4 steps by Imaris (Oxford Instruments): 1) spot detection with background correction; 2) tracking; 3) track filtering; 4) track statistics. The tracks that lasted >75% of the total time points were included in further analysis. The fluorescence intensity of the NiV-M puncta at individual time points in each track was extracted and plotted as a curve to reflect the dynamics of intensity change of a NiV-M assembly site. Several hundreds of fluorescence intensity curves obtained from NiV-M tracks in PK13 cells were used to identify a reference pattern of the NiV-M fluorescence dynamics according to their distance to the core in the t-SNE (t-distributed stochastic neighbor embedding) plot. The t-SNE plot was generated as previously described (26). This reference pattern was used to select biologically significant tracks in all treatment groups according to their fluorescence intensity curves. This selection was performed by calculating the correlation coefficient of the intensity curve of a track with the reference, and tracks with a correlation coefficient ranked in the top 20% were included in further analysis. The fluorescence curves of the selected tracks were averaged to obtain an overall pattern of the fluorescence dynamics of each treatment group. The phase identification and assembly rate determination were performed using custom-written Python codes available on GitHub(https://github.com/QLlab). Briefly, the phase was identified by determining the inflection points of the intensity curve by using a polynomial fitting shown in Equation (1):

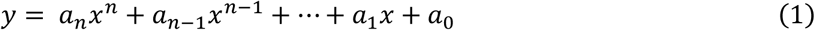

where *a*_*n*_ are coefficients, *y* is the fluorescence intensity of NiV-M puncta, *x* is the time points, *n* is the degree of polynomial optimized to 3 in our dataset. The polynomial fitting curve exhibited a snake-like trend, characterized by an initial increase, followed by a plateau, and then a decrease. The plateau (phase II) is the region between the two infection points, representing the membrane-dwelling stage. The inflection points that mark the shoulder of the curve were determined by calculating second derivatives. Phase I is the initial increase region ending at the first inflection point, representing the assembly stage. We determined the assembly rate by averaging the derivatives at the first, second, third, and fourth quartiles of phase I. Phase III begins at the second inflection point and continues to the end, representing the virus release stage.

Similarly, the fluorescence intensity of diffused GFP-tagged NiV-M constructs was determined using the zeiss TIRF microscopy at epi-illumination in live cell chamber. Images were acquired at a 1 hr interval over 18 hrs or at 5 s interval over 45 minutes, beginning at 6 hrs post-transfection of PK13 cells with GFP-NiV-M. To monitor the dynamic changes of membrane protrusions induced by NiV-M, PK13 cells transfected by GFP-NiV-M were treated by 200 μM CK-666 treatment at 6 hrs post-transfection and subjected to TIRF-M immediately. Images were acquired at a 1-hr interval over 18 hrs. Puncta and finger-like structures in each image were identified using Imaris. The ratio of the number of finger-like structures to that of puncta was calculated for each time point and normalized to the ratio at time 0 (6 hrs post-transfection).

### Cell viability assay

Cell viability was assessed using the Cell Counting Kit 8 (WST-8/CCK8) (Abcam, ab228554). Briefly, PK13 cells were seeded at a density of 6,000 cells per well in 96-well plates (Costar) and allowed to adhere overnight at 37°C. On the following day, the medium was replaced with 100 μl of fresh culture medium containing varying concentrations of CK-666. Cells were then incubated at 37°C for the specified time. After incubation, 10 μl of WST-8 reagent was added to each well, and plates were further incubated at 37°C for 1 hr. Fluorescence was measured at 460 nm using a TECAN Spark 10M fluorescent plate reader to quantify cell viability.

### Statistical analysis

Statistical analyses were performed using GraphPad Prism. The sample size, number of biological replicates, and statistical tests used for each analysis are specified in the figure legends.

## Acknowledgements

We thank the multiscale imaging facility for instrument use. We thank Dr. Youssef Chebli at McGill University for the support on laser scanning confocal microscopy and TIRF microscopy.

## Author contribution

Conceptualization: J.W. and Q.L.; Image acquisition: J.W., V.K., Q.L.; Data analysis and code writing: J.W., V.K., J.L., and Q.L.; Functional analysis: J.W., V.K., G.L.M., Q.W., Y.L., M.Z. and G.L.; Resources: Q.L.; Supervision: G.L. and Q.L.; Funding Acquisition: Q.L.. Writing: original draft: J.W., J.L., and Q.L.; revision: all authors.

## Code, reagents, and data availability

Custom-written codes are deposited in GitHub. Reagents are provided by the corresponding author on a completed material transfer agreement. Requests for the reagents and data should be submitted to.

---

**Fig. S1-related to Fig. 1.**
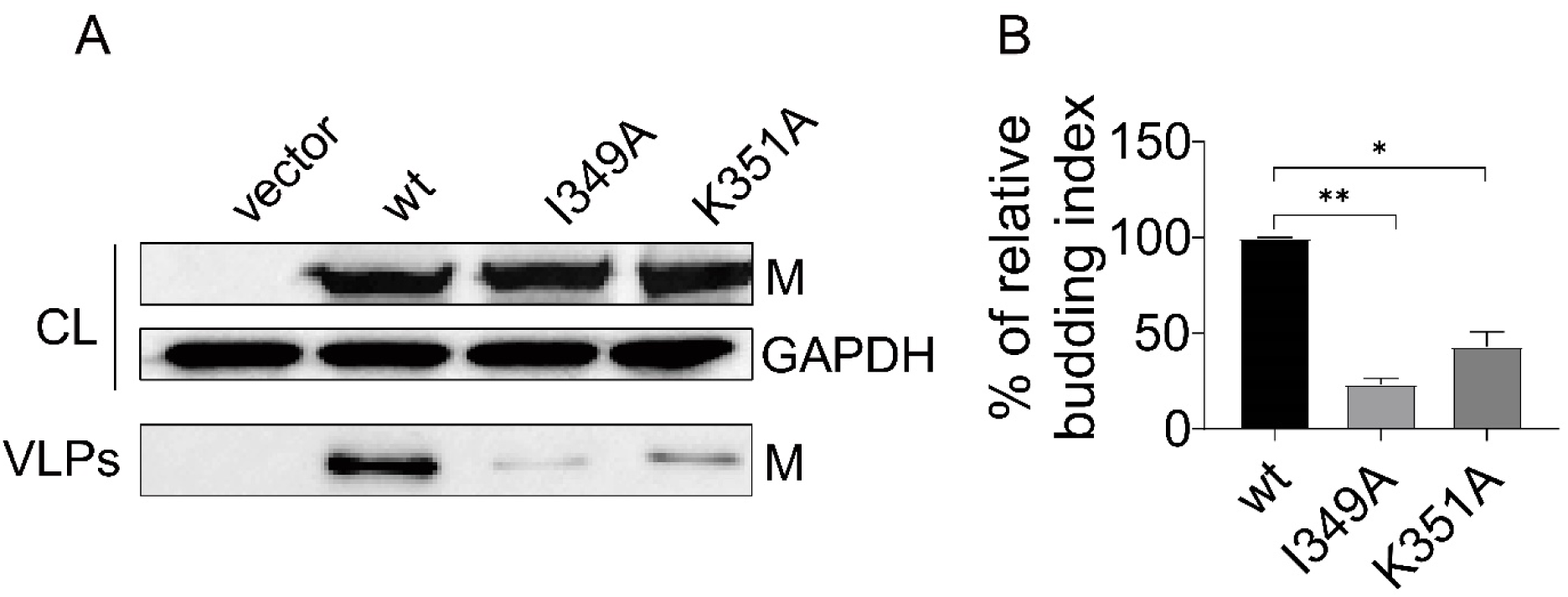
VLP production of NiV-M constructs fused with N-terminal GFP tag. (**A**) VLPs and cell lysates harvested from 293T cells transfected with empty pcDNA3 vector (vector) and plasmids coding for GFP-tagged wt, I349A, and K351A were analyzed using western blot. GAPDH was used as a loading control. NiV-M was detected using a goat anti-GFP antibody and mouse anti-GAPDH antibody. Proteins were detected by HRP-conjugated secondary antibodies. (**B**) Relative budding index is determined based on integrated immunoblot density. Results from at least three independent experiments are shown. Bars represent mean ± SEM. *p* value was obtained using Welch’s *t*-test. *: *p* ≤ 0.05, **: *p* ≤ 0.01.

**Fig. S2-related to Fig. 2.**
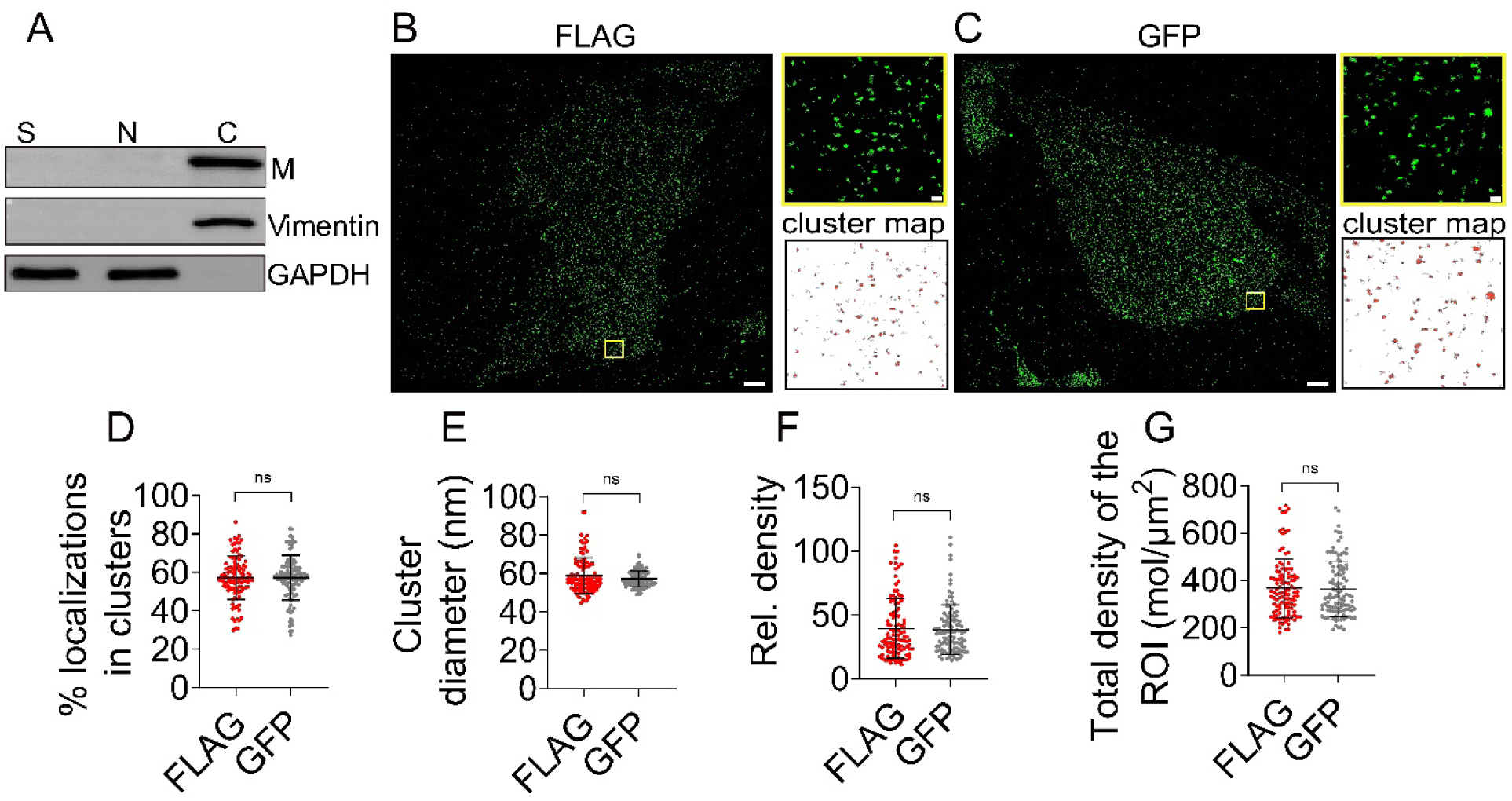
NiV-M is associated with the host cytoskeleton and its clustering pattern remains similar regardless of the labeling tags. (**A**) NiV-M is associated with the cytoskeleton in PK13 cells. PK13 cells were transfected with 3×FLAG-NiV-M for 48 hrs and the soluble compartment (S), nuclear compartment (N), and cytoskeleton (C) were extracted. The partitioning of NiV-M to each compartment was analyzed by western blot analysis. GAPDH is the control for S and N compartments, and vimentin C compartment. NiV-M was detected using a mouse anti-FLAG antibody, GAPDH a mouse anti-GAPDH antibody, and vimentin a rabbit anti-vimentin antibody. (**B-G**) The effect of 3×FLAG and GFP tags on NiV-M clustering. PK13 cells expressing 3×FLAG-NiV-M or GFP-NiV-M were subjected to SMLM imaging. (**B, C**) x-y cross-section (600 nm thick in z) of the SMLM images of 3×FLAG-NiV-M (**B**) and GFP-NiV-M (**C**) at the ventral membrane of PK13 cells. The boxed regions are enlarged, and the cluster maps of the NiV-M localizations are shown. Unclustered NiV-M localizations are gray, while clustered NiV-M localizations are red in the cluster maps. Scale bar: 1 μm and 200 nm. The percentage of localizations in clusters (**D**), cluster diameter (**E**), relative density (**F**), and total density of the ROI (mol/μm^2^) (**G**) are shown in dot plots. Sample size n = 108 (3×FLAG-NiV-M) and 108 (GFP-NiV-M) from 10-20 cells per group. Bars represent mean ± SD. *p* value was obtained using Welch’s *t*-test. ns: *p* > 0.05.

**Fig. S3-related to Fig. 5.**
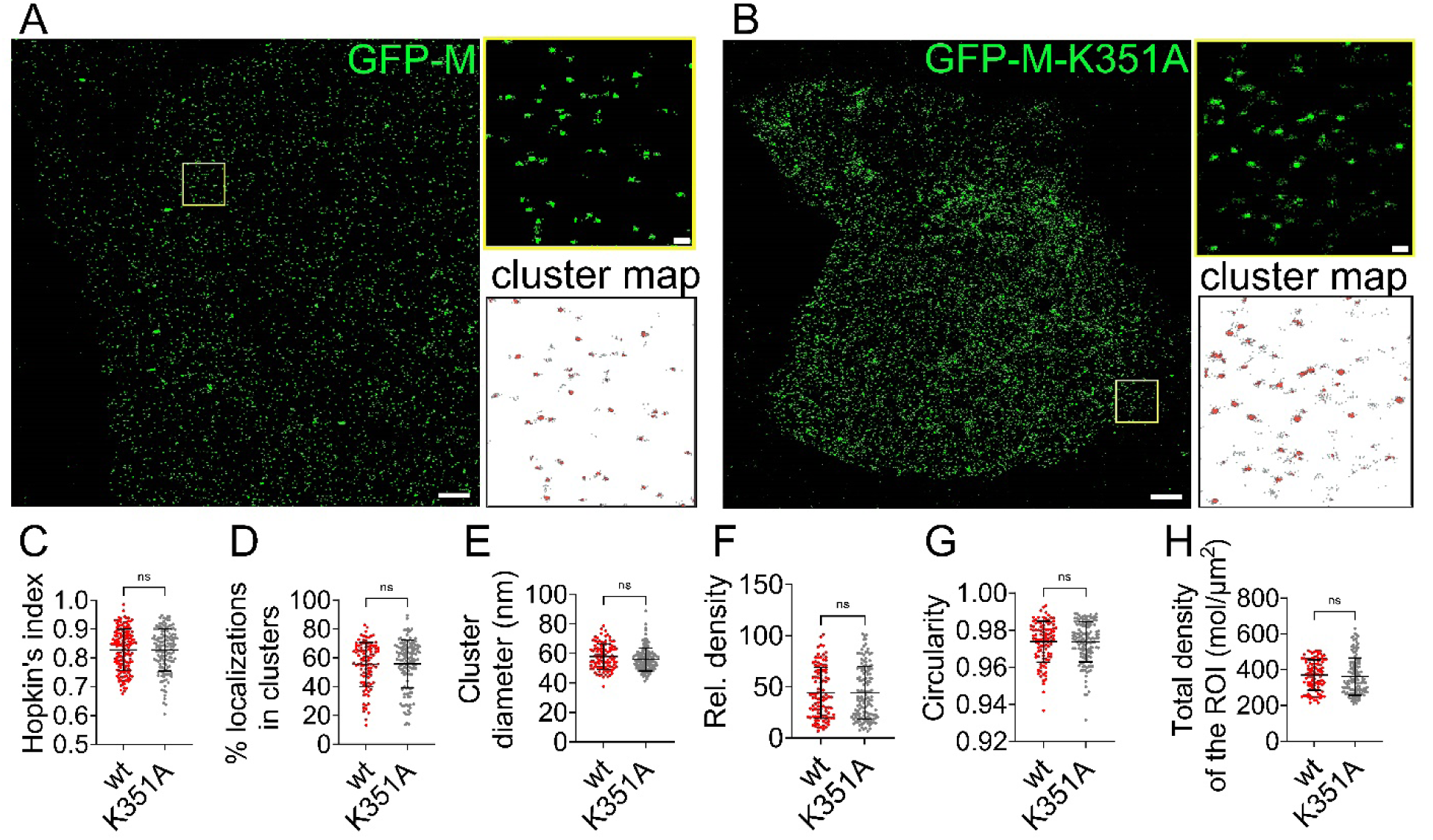
The K351A mutant at the actin-binding domain did not alter the nano-organization of NiV-M. PK13 cells overexpressing GFP-tagged NiV-M-wt and K351A were subjected to SMLM imaging. (**A** and **B**) x-y cross-section (600 nm thick in z) of the SMLM images of GFP-tagged NiV-M (**A**) and K351A (**B**) at the ventral membrane of PK13 cells. The boxed regions are enlarged, and the cluster maps of the NiV-M localizations are shown. Scale bars: 1 μm and 200 nm. (**C**) The Hopkin’s index of the localizations of NiV-M-wt and K351A. (**D-H**) The percentage of localizations in clusters (**D**), cluster diameter (**E**), relative density (**F**), circularity (**G**), and total density of the ROI (**H**) were plotted. Sample size n = 105 (wt) and 140 (I349A) from 12-18 cells per group. Bars represent mean ± SD. *p* value was obtained using Welch’s *t*-test. ns: *p* > 0.05.

**Fig. S4-related to Fig. 6.**
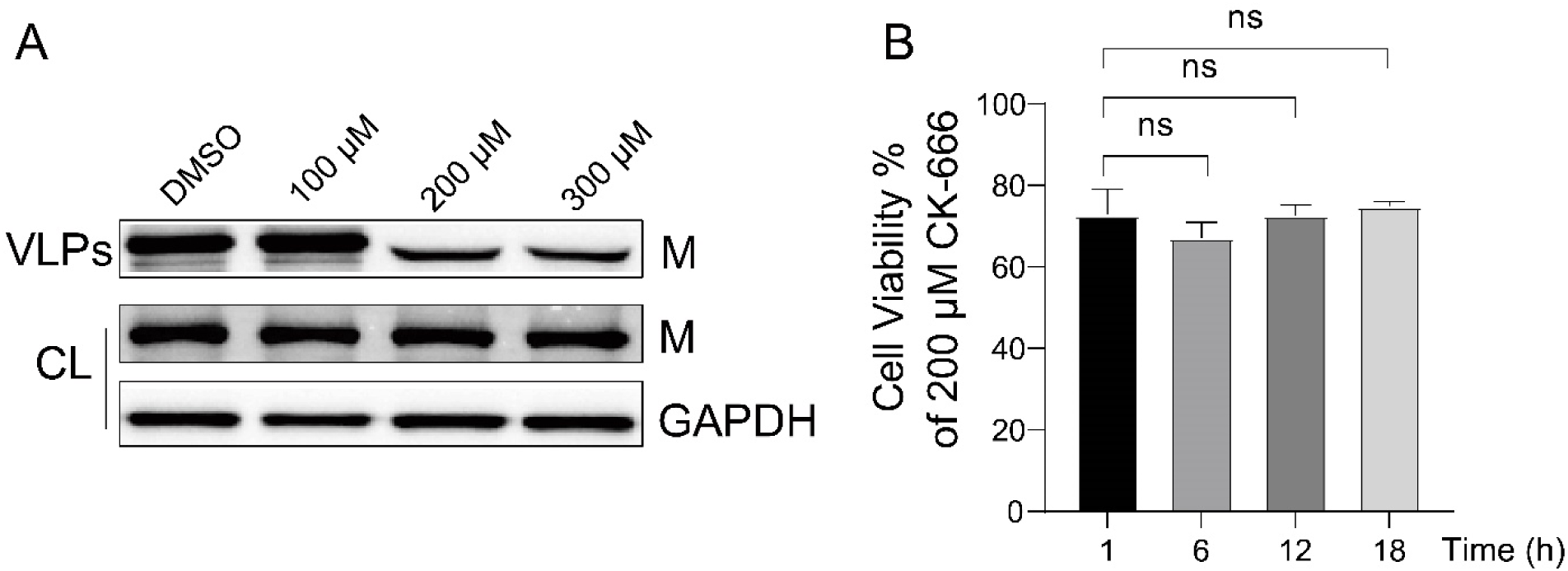
The effects of CK-666 on NiV VLP production and PK13 cell viability. (**A**) VLPs and cell lysates were harvested from PK13 cells transfected with 3×FLAG-NiV-M and treated with 300 μM DMSO, 100 μM CK-666, 200 μM CK-666 and 300 μM CK-666. NiV-M in the cell lysate (CL) and VLPs were analyzed using Western Blot. (**B**) PK13 cells were treated with 200 μM CK-666 for 1 h, 6 h, 12 h, and 18 h, respectively. The cell viability was detected using a CCK8 kit. Bars represent mean ± SD. *p* value was obtained using Welch’s t-test. ns: *p* > 0.05.

